# Formation of Condition-Dependent Alpha-Synuclein Fibril Strain in Artificial Cerebrospinal Fluid

**DOI:** 10.1101/2025.02.21.639308

**Authors:** Rūta Sniečkutė, Darius Šulskis, Arūnė Jocytė, Urtė Venclovaitė, Rimgailė Tamulytė, Mantas Žiaunys, Vytautas Smirnovas, Andrius Sakalauskas

## Abstract

α-Synuclein (aSyn) is an intrinsically disordered protein involved in neurotransmission and synaptic plasticity. The pathological aggregation of this protein is a hallmark of synucleinopathies such as Parkinson’s disease (PD) or Multiple System Atrophy (MSA). Misfolded aSyn, which primarily originates in cell cytosol, transmits between neurons, promoting a prion-like propagation. However, the extracellular environments such as interstitial and cerebrospinal fluids (ISF & CSF) play a major role in its clearance and pathological transformation. The molecular components of CSF, including proteins, glycosaminoglycans, and metal ions may influence the aggregate morphology, structure and cytotoxicity to cells. To better understand how extracellular composition affects aggregates and their formation, we employed artificial cerebrospinal fluid (aCSF) to mimic potential aggregation processes occurring in CSF. We observed distinct aCSF-specific aSyn fibrils that exhibited low stability outside aCSF, and the removal of key CSF components led to its structural alterations. Cryo-electron microscopy revealed that these fibrils possess an electron density pocket coordinated with polar basic AAs (K43, K45, H50) that is also observed in aggregates obtained from MSA patients. Our findings illustrate the importance of physiologically relevant conditions in studying aSyn aggregation and may explain why disease-related fibril structure replication *in vitro* has not yet been successful.

## Introduction

α-Synuclein (aSyn) is a protein primarily found in the brain, playing a crucial role in neuronal functions such as synaptic vesicle regulation and neurotransmitter release^1,2^. Its abnormal aggregation is a hallmark of several neurodegenerative diseases, including Parkinson’s disease (PD), dementia with Lewy bodies and other synucleinopathies^3,4^. In these pathologies, aSyn misfolds and forms insoluble aggregates in neurons that are called Lewy bodies and Lewy neurites^5^. Accumulation of abnormal aggregates leads to impaired neurotransmitter release, mitochondrial dysfunction, loss of presynaptic proteins, and disruption of protein trafficking that generally results in neuronal loss and neurodegeneration^6^.

One of the key aspects of aSyn is its ability to propagate between cells via different pathways^7,8^ and in diverse pathological forms – soluble oligomers and insoluble aggregates. aSyn and its soluble misfolded species are mostly contained within the cell, however, there is evidence that it may spread by diffusion to the intercellular space^9^. This can occur via (1) direct translocation across the membrane that is facilitated by specific protein such as DnaJ homolog subfamily C member 5 (DNAJC5)^10^, (2) by calcium-mediated exosomal secretion^11,12^ or other neuronal activity-dependent pathways^13^. Soluble aSyn species that escaped the cell can be absorbed by other neurons mostly through endocytosis and passive diffusion across the plasma membrane^14,15^. However, aSyn aggregates formed within the cell can be secreted from the neuron, enabling the interaction with the neighbouring neurons or immune cells which can uptake them. This process contributes to spreading the pathology across brain regions following a prion-like mechanism^16,17^. The aggregate transmission may occur through exocytosis and endocytosis, or through tunnelling nanotubes for larger assemblies^18^. Once inside the recipient neuron, the misfolded aSyn induces native aSyn to misfold and aggregate, perpetuating a cycle of toxic accumulation leading to cell death and progression of neurodegeneration^19,20^.

While aSyn aggregation is considered to originate within the neuron at presynaptic terminals^21,22^, different transport pathways of this protein and its misfolded species through synapses play a crucial role in disease development. The neuron environment consisting of interstitial and cerebrospinal fluids (ISF & CSF) is crucial for maintaining the brain’s homeostasis and ensuring proper neuronal function^23,24^. These fluids are part of the brain’s microenvironment, and their interaction is important for waste removal, delivery of nutrients, and overall brain health^25^. In the case of disease pathology, the extracellular space contains various forms of the protein, from native aSyn to toxic misfolded species^26^. Here, the constant exchange between ISF and CSF plays a major role in clearing aSyn from the brain^27^, highlighting the importance of their interaction in the context of neurodegenerative diseases^28^.

The misfolded aSyn deposits within the extracellular space may change, seed the aggregation of cell-free aSyn^29^ and result in various fibril conformations due to shifted environment conditions^30,31^. Differences in crowding, ionic strength and reaction component composition play a crucial role in formation of both *de novo* and seeded aggregates^32,33^. There is evidence that glycosaminoglycans (such as heparin)^34^, metal ions^32^, lipoproteins^35^ or other proteins such as human serum albumin^36^ shape the formation of aSyn fibrils resulting in different aggregate structures. This is why, it is no surprise that multiple distinct aSyn aggregate structures formed *in vitro* and *in vivo* were recorded using cryo-electron microscopy^37–40^, with cases describing unknown molecules within the core of the aggregate^41,42^. Such residual molecules from either cellular or extracellular environments suggest the importance of studying the aggregation of aSyn under physiologically relevant conditions.

The study of aSyn fibril formation in the extracellular environment may provide critical insights into the development of aSyn related neurodegenerative diseases. While it was not yet possible to template the disease-derived fibril structure using recombinant aSyn protein under *in vitro* conditions^41^, the search for a model system that would help in simplifying the templating process and improving our understanding of disease-related mechanism is very important. In this research, we have used artificial cerebrospinal fluid (aCSF) that contains major CSF components to simulate the aggregation of aSyn in the extracellular environment. The aCSF conditions yielded a distinct aggregate conformation with a low stability outside the aCSF environment, while removal of major cerebrospinal fluid components resulted in structural variation between the formed aggregates. Cryo-electron microscopy imaging analysis showed a three-dimensional structure with a small molecule entrapped within the fibril core. These results demonstrate that mimicking the CSF environment is essential for retaining aSyn aggregate structures formed within aCSF, which may lead to successful replication of *ex vivo* aggregates *in vitro*.

## Materials and methods

### Preparation of aCSF

The preparation of artificial cerebrospinal fluid (aCSF) was described previously^43^. The resulting composition of aCSF contains the main components that may have an effect on protein aggregation process. The compounds containing Na^+^ and K^+^ ions were dissolved as one stock solution called phosphate buffer (further referred as PB (A component)). Meanwhile, the rest of the components were dissolved separately (B-I) as described (Supplementary table 1). HCl was added to the final 1x aCSF solution to yield a pH of 7.33.

### Expression and Purification of aSyn

Recombinant aSyn was expressed and purified as described in previous work^44^. In short, *E.coli* BL(21) Star (DE3) as host cells were transformed with pDS5 plasmid containing SNCA gene. Prepared cells were grown overnight at 37°C, in ZYM-5052 medium supplemented with 100 µg/ml ampicillin. Harvested cells were pelleted, resuspended in lysis buffer and homogenized. Later, the collected supernatant was heat treated at 80°C for 20 min in a water bath. The resulting suspension was centrifugated. Ammonium sulphate was added to the collected supernatant (to saturation of 42%) to precipitate the target protein. After centrifugation the pellet was dissolved and dialyzed. The final solution containing aSyn and 20 mM Tris-HCl (pH 8.0) (supplemented with 0.5 mM DTT and 1 mM EDTA) buffer solution was loaded onto DEAE sepharose sorbent for ion exchange chromatography. The target protein was eluted by using 20 mM Tris-HCl (pH 8.0), 1 M NaCl, 0.5 mM DTT, 1 mM EDTA buffer solution. Collected fractions were combined, concentrated and loaded onto prepacked Hi-Load 26/600 Superdex 75 pg. size exclusion column that was equilibrated in PB (127 mM NaCl, 1.8 mM KCl, 7.81 mM Na_2_HPO_4_, 3.19 mM NaH_2_PO_4_, 1.2 mM KH_2_PO_4_, pH 7.33). aSyn was eluted at 130 – 155 mL, collected and adjusted with elution buffer to final 100 µM concentration. aSyn solution was distributed into tubes (1.0 mL volume) and stored at −20°C. The prepared aSyn stock solution aliquots were used a single time after thawing, to avoid any potential oligomerization process during re-freezing conditions.

### Aggregation Experiments

The *de novo* aggregation of aSyn was performed under various conditions as described (Supplementary table 2). Purified 100 µM aSyn in PB (127 mM NaCl, 1.8 mM KCl, 7.81 mM Na_2_HPO_4_, 3.19 mM NaH_2_PO_4_, 1.2 mM KH_2_PO_4_, pH 7.33) was mixed with 10xPB and 20x-100x B - I stock solutions (aCSF components), 10 mM Thioflavin T (ThT) stock solution and MilliQ water to yield a final aSyn concentration of 50 µM and 100 µM ThT in final reaction mixture. The prepared final solutions were distributed into Greiner non-binding 96-well plates with 80 µL/well and each well containing a 3 mm glass-bead. Aggregation kinetics were monitored by using Clariostar Plus plate reader with the following parameters: 37 °C, orbital agitation of 600 RPM, excitation and emission wavelengths of 440 nm and 480 nm, ThT fluorescence measured every 5 minutes.

The seeded aggregation experiment was conducted by preparing a reaction mixture identical to that used for *de novo* aggregation, with the addition of 10% of an aggregated aSyn sample. Each individual repeat was distributed across four wells of the 96-well plate. Aggregation was monitored using the same aggregation protocol as for *de novo* aggregation process. For seeded aggregation of where *de novo* prepared aCSF fibrils were used to initiate the aggregation of aSyn in aCSF and PB solutions, the process was monitored without agitation. The resulting aggregate solutions were placed in non-binding tubes (Axygen, maximum recovery, ref. MCT-200-L-C) and centrifuged for 10 min at 15,000 x g. The supernatant was used for measuring the unaggregated (soluble) fraction of aSyn in each sample.

### Fibril restructurization measurements

The aCSF aSyn sample was centrifugated for 10 min at 15 000 x g, after which the 90 % of supernatant volume was removed. The concentrated fibril solution was placed in Corning non-binding 96-well plates (15 µL solution and one 3 mm glass bead in each well). The wells were then supplemented with either 135 µL aCSF or PB solutions (each containing 50 µM ThT), followed by brief pipetting. Immediately after the mixing procedure, the sample excitation-emission matrices (EEM) were scanned as described previously^45^. The scanning procedure was repeated every 60 seconds for 30 min with 5 second 600 RPM agitation before each measurement. EEM intensity maximum positions and values were determined as described previously^45^.

### Aggregate resuspension in Buffer A and different aCSF composition

100 µL of the final aggregate solution was placed in 250 µL PCR tubes that were centrifugated 5 minutes at 9000 RPM (ThermoFisher Mini Centrifuge). Then, the supernatant was removed and the fibrils were resuspended with 100 µL of Buffer A or different aCSF composition solution. This process was repeated once again before imaging of the sample was conducted.

### Fourier-Transform Infrared Spectroscopy (FTIR)

The aSyn fibril solutions were centrifuged for 10 min at 15,000 x g. The supernatant was removed, and the remaining fibril pellets were resuspended in 200 µL of D_2_O (containing 300 mM NaCl)^46^. The centrifugation and resuspension into D_2_O were repeated three times. After the final centrifugation the supernatant was removed, and the remaining sample was resuspended in 80 µL of D_2_O with 300 mM NaCl. The measurements were conducted as described previously^45^. All data processing was performed using QUASAR software^47^.

### Atomic Force Microscopy (AFM)

aSyn fibrils formed in Buffer A and aCSF were used to prepare samples for AFM imaging similarly as previously described^43^. In short, freshly cleaved mica was functionalized by placing 40 µL of 0.5 % (v/v) APTES (Sigma-Aldrich, cat. No. 440140) in MilliQ water and incubating for five minutes. Then the mica was washed with 2 mL MilliQ water and dried under gentle airflow. 40 µL of each diluted sample (diluted to 20 µM aSyn concentration) was added to individual mica, incubated for 5 minutes, washed and dried as previously described. The AFM imaging was done using Dimension Icon (Brucker) atomic force microscope. Images of 1024 x 1024 pixel resolution were recorded using Nanoscope 10.0 (Bruker) operating in tapping mode and equipped with silicon tip (Tap300AI-G; 40 Nm^−1^, Budget Sensors). Image levelling, corrections, cropping and analysis was done using Gwyddion 2.63 software^48^. The fibril profile heatmaps were drawn using Origin 2018. Gaussian fitting was done in order to approximate fibrils height and width traces, which were used to calculate Max-Min and Pitch parameters.

### High-Speed Atomic Force Microscopy (HS-AFM)

aSyn fibrils formed in aCSF were placed on the APTES-functionalized mica and incubated for 5 minutes. Then the mica was washed with aCSF solution and was placed in a custom measurement cell (100 µL) with aCSF or PB solutions. The imaging was done using HS-AFM (SS-NEX, RIBM, Japan). Images of 250 x 250 pixel resolution were recorded using IgorPro-based RIBM software (Ibis v.1.0.2.4, IgorPro v.6.3.7.2). The system operated in tapping mode and was equipped with ultrashort (8 µm) silicon nitride (Si3N4) rectangular cantilevers (BL-AC10DS-A2, Olympus, Japan) with a tip radius of 24 nm. These cantilevers had a nominal spring constant of approximately 0.1 N/m and a resonant frequency of approximately 0.5 MHz in solution. Data analysis was performed using NanoLocz (v.1.20) software^49^.

### Cryo-Electron Microscopy (Cryo-EM) data collection

For Cryo-EM sample preparation, 3 μL of alpha synuclein fibrils were applied to the glow-discharged holey carbon Cu grids (Quantifoil) and blotted with filter paper using Vitrobot Mark IV (FEI Company). The grids were immediately plunge-frozen in liquid ethane and clipped. Cryo-EM data was collected either on Glacios transmission electron microscope (Fisher Scientific) operated at 200 kV camera or Krios transmission electron microscope (Fisher Scientific) operated at 300 kV and equipped with Falcon 3EC/4i cameras. Cryo-EM data collection statistics can be found in Supplementary Table 3.

### Image pre-processing and helical reconstruction

The micrographs were aligned, motion corrected using MotionCorr2 1.2.1^50^ and the contrast transfer function was estimated by CTFFIND4^51^. The fibrils were picked and all subsequent 2D classifications were performed in Relion 5.0^52^. Distribution of polymorphs was identified by FilamentTools (https://github.com/dbli2000/FilamentTools) as a part of Relion software and analysis dendrograms are presented in Suplementary Figures 1-3. Segments with box size of 1024 downscaled to 384 were used to generate de novo model using relion_helix_inimodel2d function. Next, three-dimensional auto-refinements were performed with optimisation of the helical twist and rise parameters once the resolutions extended beyond 4.7 Å. To improve the resolution, Bayesian polishing and CTF refinement were perfomed. Final maps were sharpened using standard post-processing procedures in RELION (Supplementary figure 4). Helical model parameters can be found in Supplementary Table 3.

### Model building and refinement

The atomic model was built de novo in Coot^53^ and was subjected to several real-time refinements in PHENIX^54^ with manual curation of outlines to ensure energy-favoured geometry. Map volume and models were visualized using Chimerax program^55^. Hydrophilicity and hydrophobicity of model was inspected with ProCart webserver (https://github.com/jianglab/procart)^56^. Validation report is attached as a Supplementary document I.

### Cell culturing

Cell culture used for experiments (SH-SY5Y human neuroblastoma cells) was obtained from the American Type Culture Collection (ATCC, Manassas, VA, USA). The cells were grown in Dulbecco’s Modified Eagle Medium (DMEM) (Gibco), supplemented with 10% Fetal Bovine Serum (FBS) (Sigma-Aldrich), 1% Penicillin–Streptomycin (10,000 U/mL) (Gibco) at 37 °C in a humidified, 5% CO_2_ atmosphere in a CO_2_ incubator.

### MTT and LDH assay

The samples for cell assays were collected from seeded aggregation experiments. To avoid any possible effect of reaction buffers (PB and aCSF), samples containing aSyn fibrils were centrifuged for 1 h at 15 000 x g. The supernatant was removed, and the remaining fibril pellet resuspended in the same volume of DMEM medium (for MTT assay) or in Advanced DMEM (for LDH assay).

For both cell assays SH-SY5Y cells were seeded in a 96-well plate (∼15,000 cells/well) and incubated overnight. In case of MTT assay, after the incubation cell medium was changed to the one containing aSyn fibrils formed either in PB or in aCSF. For each of the conditions titration of sample was applied to yield the final concentrations of 20, 10, 5 and 1 µM. After 48 h of incubation 10 µM of 3-(4,5-dimethylthiazol-2-yl)-2,5-diphenyltet-tetrazolium bromide (MTT) reagent (12.1 mM in PBS) was added to each well and followed by 2 h of incubation. To dissolve formazan crystals 100 µL of 10% SDS with 0.01 N HCl solution was added to each well. After 2 h, the absorbance was measured at 570 nm and 690 nm (as reference wavelength) using a Clariostar Plus plate reader.

Meanwhile, LDH release into the medium was quantified by using Cytotoxicity Detection kit (Roche). After overnight incubation, the medium inside the wells was aspirated, 100 µL/well of Advanced DMEM (Gibco) was added. Then, each sample was diluted in Advanced DMEM. Prepared samples were added to the wells to reach 200 µL/well of total medium volume, which resulted in the final 20, 10, 5 and 1 µM sample concentrations for each condition. After 24 h of incubation, 100 µL of the medium from each well was aspirated, centrifugated, and transferred into a flat bottom 96-well test plate. Freshly prepared LDH reagent was added and incubated for 30 min at room temperature. The absorbance was measured at 492 nm and 600 nm (as reference wavelength) using a Clariostar Plus plate reader.

### Cell imaging

The aSyn fibrils for cell imaging were pre-formed by using mCherry-aSyn fluorescent protein. Here the seeded aggregation experiment was conducted as described in methods section *Aggregation Experiments*, with the addition of 5% mCherry-aSyn.

The SH-SY5Y cells were seeded in a 24-well plate (∼90,000 cells/well) and incubated overnight. Samples containing mCherry-aSyn in PB and mCherry-aSyn in aCSF were centrifuged for 1 h at 15 000 x g. Pelleted fibrils of each sample were resuspended in FluoroBrite DMEM (Gibco) and further diluted to a final concentration of 20 µM or 1 µM. After the overnight incubation with samples, cells were analysed with EVOS FL Auto fluorescence microscope (Life Technologies, USA) by taking photos with 20x and 40x objectives. Here two types of data were collected: “before” and “after”. For “after” imaging, media containing aSyn samples was removed, cells washed with phosphate-buffered saline (PBS, Gibco) and replaced with new FluoroBrite media.

All images were analysed using the Fiji package for ImageJ.

### Statistical Analysis

The *de novo* aSyn aggregation kinetic data (lag time and apparent rate) was analysed by fitting the kinetic curves using Boltzmann’s sigmoidal equation^45^. The seeded aggregation kinetic curves were normalized and then analysed by linear fit. Halftime and slope parameters were extracted from the fitted data. Each condition was repeated 8 times to evaluate the reproducibility of the data.

The statistical analysis of MTT and LDH assay results was conducted by using ORIGIN 2018 software (OriginLab Corporation, MA, USA), One-way analysis of variance (ANOVA) Bonferroni means comparison (n = 9). Three independent sets of samples with three repeats of each condition were used for MTT and LDH assays.

## Results

### aCSF components stabilize the aggregation process of aSyn

To understand the influence of formulated artificial cerebrospinal fluid (aCSF) on the aSyn aggregation process, we have designed kinetic experiments in phosphate buffer (*further referred as* – PB (A component of aCSF)), aCSF and solutions with varying aCSF components (Fig. 1 A). Since aSyn is prone to polymorphism under certain sets of conditions^45^, we evaluated the molecules within aCSF that can modify or stabilize the aSyn aggregation pathway. In order to assess the variability of aSyn aggregation, eight replicates for each condition were used. Out of all ten variations displayed, the most contrastive conditions were PB, without calcium and without magnesium that resulted in more scattered *de novo* aggregation kinetic parameters (Fig. 1 B). PB environment exhibited the most variation in aggregation lag time, but the deviation in the apparent rate constant was among the lowest. Calcium played a significant role in increasing the overall aggregation speed, whereas magnesium only affected the apparent rate constant.

**Figure 1.**
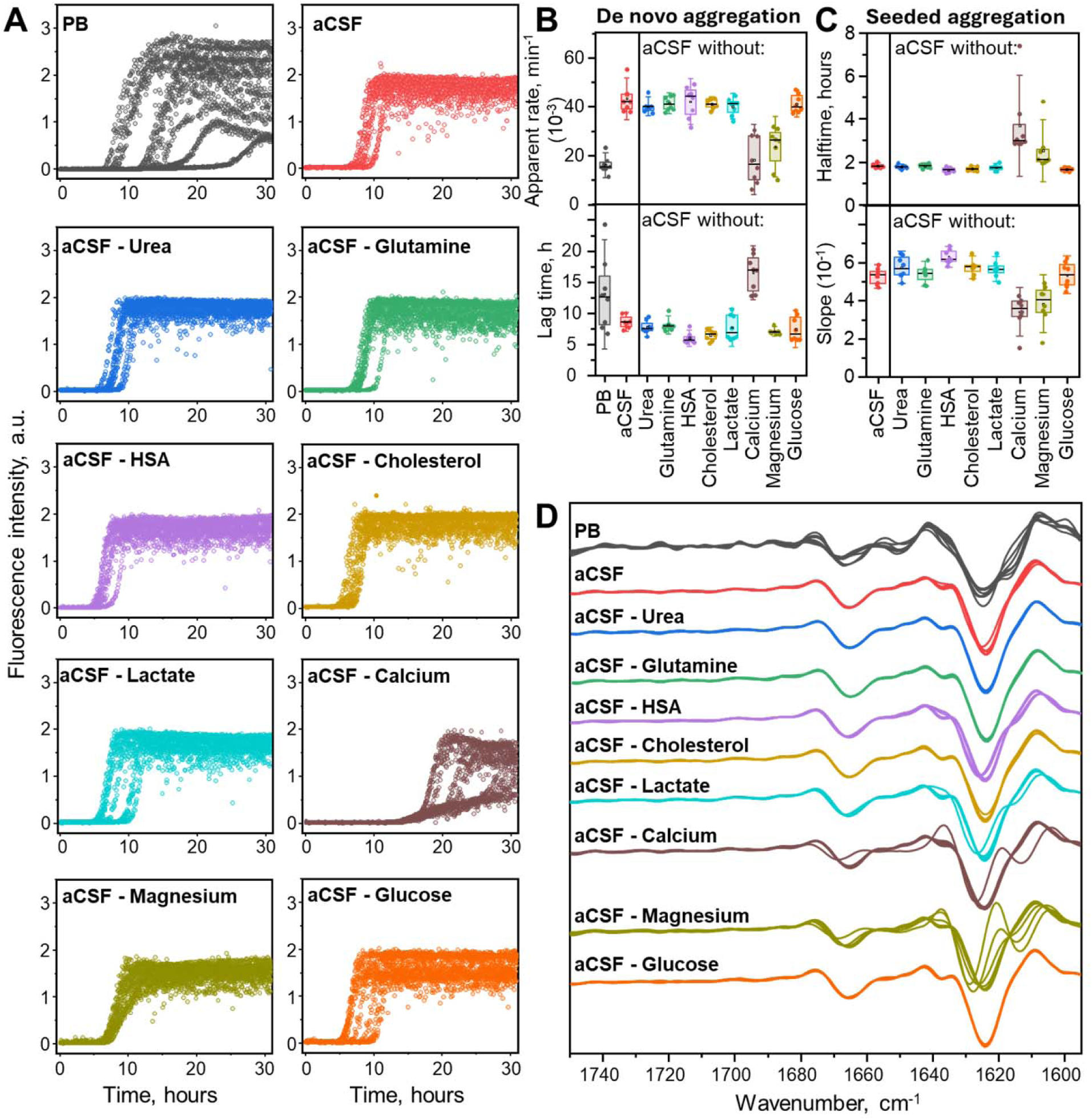
Influence of aCSF composition on aSyn aggregation kinetics and fibril secondary structure. Aggregation kinetics in different aCSF composition (A) and its parameters of *de novo* (B) and seeded (C) fibrillation process. Resulting aSyn aggregate second derivative FTIR spectra (D) are overlaid based on aCSF composition where aSyn was aggregated. Each condition is represented with separate individual repeats (n = 8).

The kinetic data variation also translated to seeded aggregation (Fig. 1 C & Supplementary Fig. 5) and the resulting fibril second derivative FTIR spectra (Fig. 1 D). The removal of calcium and magnesium resulted in slower seeded aggregation kinetics and different FTIR spectra among the technical repeats. While it is known that calcium promotes aSyn aggregation by binding to C terminal of this IDP^57^, the synergistic effect of other aCSF components (including magnesium) favour the formation of aggregates with distinct FTIR spectra second derivatives. Unlike the spectra recorded for most of the aggregates formed in aCSF conditions (including its variations), PB aggregate FTIR second derivatives showed an additional prominent minimum at 1616 – 1618 cm^−1^, indicating a stronger β-sheet region. The removal of Ca^2+^ or Mg^2+^ ions resulted in a few specific aggregate spectra exhibiting minima at 1615 – 1613 cm^−1^, and 1627 – 1629 cm^−1^, which may also correspond to formation of an aSyn aggregate type with different beta-sheet regions and stabilized orientation^58^.

### aCSF fibrils are more homogeneous than those produced in PB

To investigate the differences between the end-products of aSyn aggregation in PB and aCSF, AFM and Cryo-EM imaging were employed (Fig. 2). Due to the large quantity of samples after kinetic experiments, AFM images were taken from the samples that had distinct FTIR spectra among its group. Fibrils found in PB appeared longer and more widely dispersed than those displayed in aCSF (Fig. 2 A, Supplementary Fig. 6). Additionally, in PB multiple distinct species of fibrils were observed (Fig. 2 B). These fibrils possessed different pitch and periodicity height variation (Max-Min). PB1 and PB2 had no significant pitch difference but contrasted by periodicity height. PB3 fibrils displayed little to no periodicity or low variations, potentially due to lower AFM resolution that was used. In contrast, aCSF fibrils were clustered, reducing the proficiency of such evaluation, however, all selected fibrils were uniform in pitch and Max-Min of the periodicity. To reduce the effect of AFM sample preparation (APTES-functionalized mica and sample wash procedures), cryogenic electron microscopy was used (Fig. 2 C & Supplementary Fig. 1 – 2). The EM images confirmed similar tendencies as observed in AFM. The results revealed three distinct fibril species in PB, compared to one dominant type in aCSF. Additionally, PB type 1 had twist Max-Min difference of around 12 Å.

**Figure 2.**
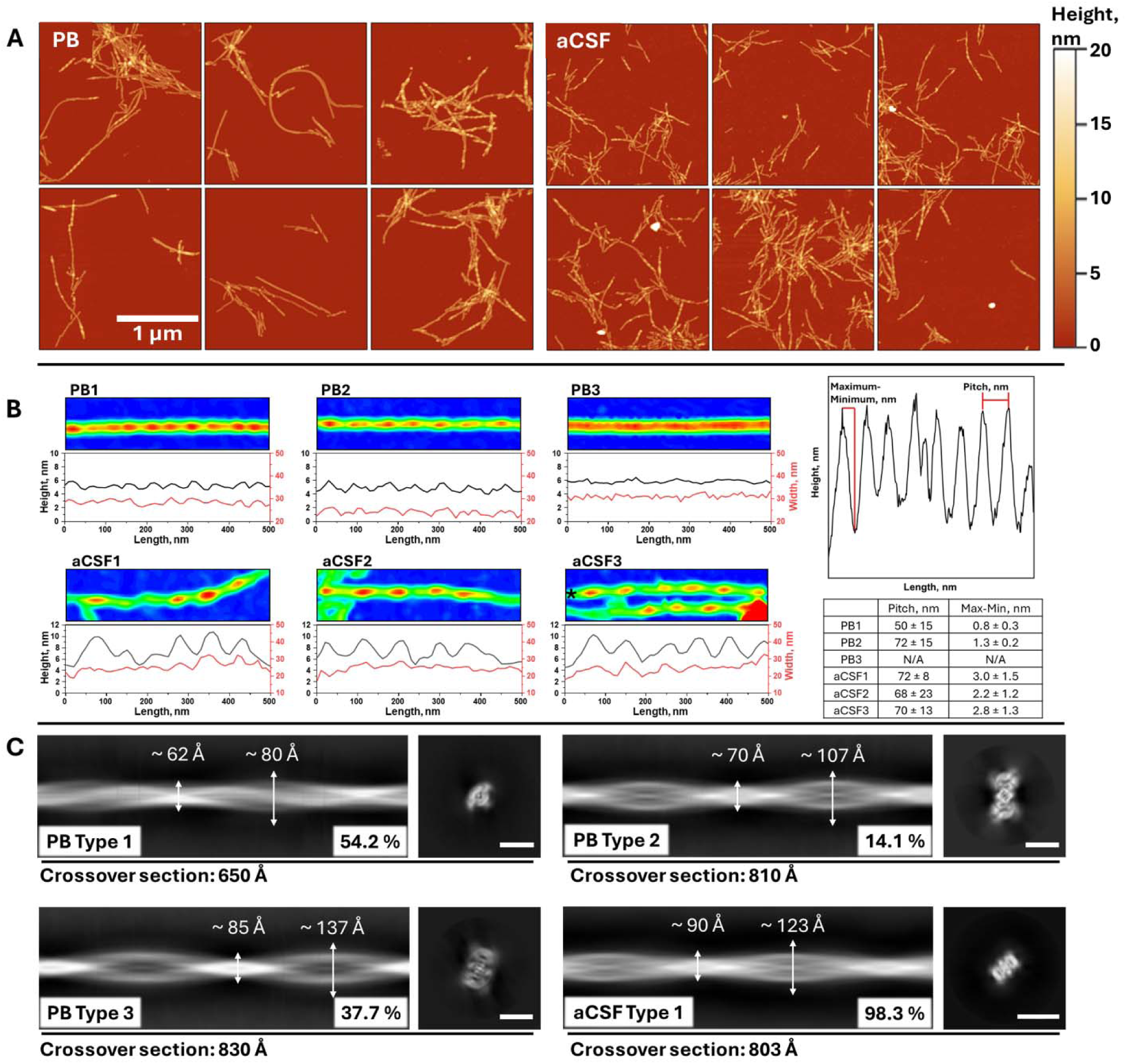
End products of aSyn aggregation recorded using AFM (A) and fibril morphology examples (B) that were formed in PB or aCSF. For each example AFM image height heatmap and its height and width values are displayed. Pitch and Max-Min averages across the fibril are summarized in the adjacent table. Distinct fibril types observed in PB and aCSF using Cryo-EM (C) along with their probability and crossover section. Scale bar represents 100 Å.

### aSyn fibrils form a unique structure in aCSF

Cryo-EM was used to solve the structure of aSyn fibrils formed in aCSF. The fibrils contained two intertwined protofilaments that exhibited a C2 symmetry with a pitch of 803 Å, helical twist – 1.10° and a helical rise of 4.91 Å (Fig. 3 A, B). During the map building, pseudo-2_1_-screw axis symmetry was also tested, but it led to a lower resolution map compared to C2, and β-sheets did not separate. The 3D reconstructed map extended from 1 to 99 residues, covering both N-terminal and entire non-amyloid beta component (NAC) regions, with the estimated final resolution of 2.8Å (Fig. 3 C). One salt bridge between lysine 21 and glutamine 35 was identified via ProteinTools webserver^59^ (Fig. 3 D) and ligand embedded within the fibril core (Fig. 3 E). The ligand binding pocket was coordinated between K43, K45, and H50 residues, with potential glucose, glutamine, various metal-ion or salt-water complexes from aCSF fitting the electron density. Similar pocket and electron densities were previously observed in filaments extracted from patients with multiple system atrophy (MSA)(Supplementary figure 7)^60^. Twelve parallel β-sheets were observed in the secondary structure, with β5 and β10 forming steric zipper motifs (Fig. 3 F). We also analyzed the distribution of charges and hydrophilicity in the solved structure (Fig. 3 G). The positive charges were clustered in the ligand-binding core and along the fibrils side, arranged by K10, K12, K58 and K60 lysines, which could allow the binding of negative charge molecules like DNA. Hydrophilic sites were also located at the same positions, whereas most hydrophobic space was between β1 and β6 sheets. The overall structural isomorph did not resemble previously solved structures in CSF^26^, which indicates aggregation in aCSF conditions allows the formation of a unique aSyn conformation.

**Figure 3.**
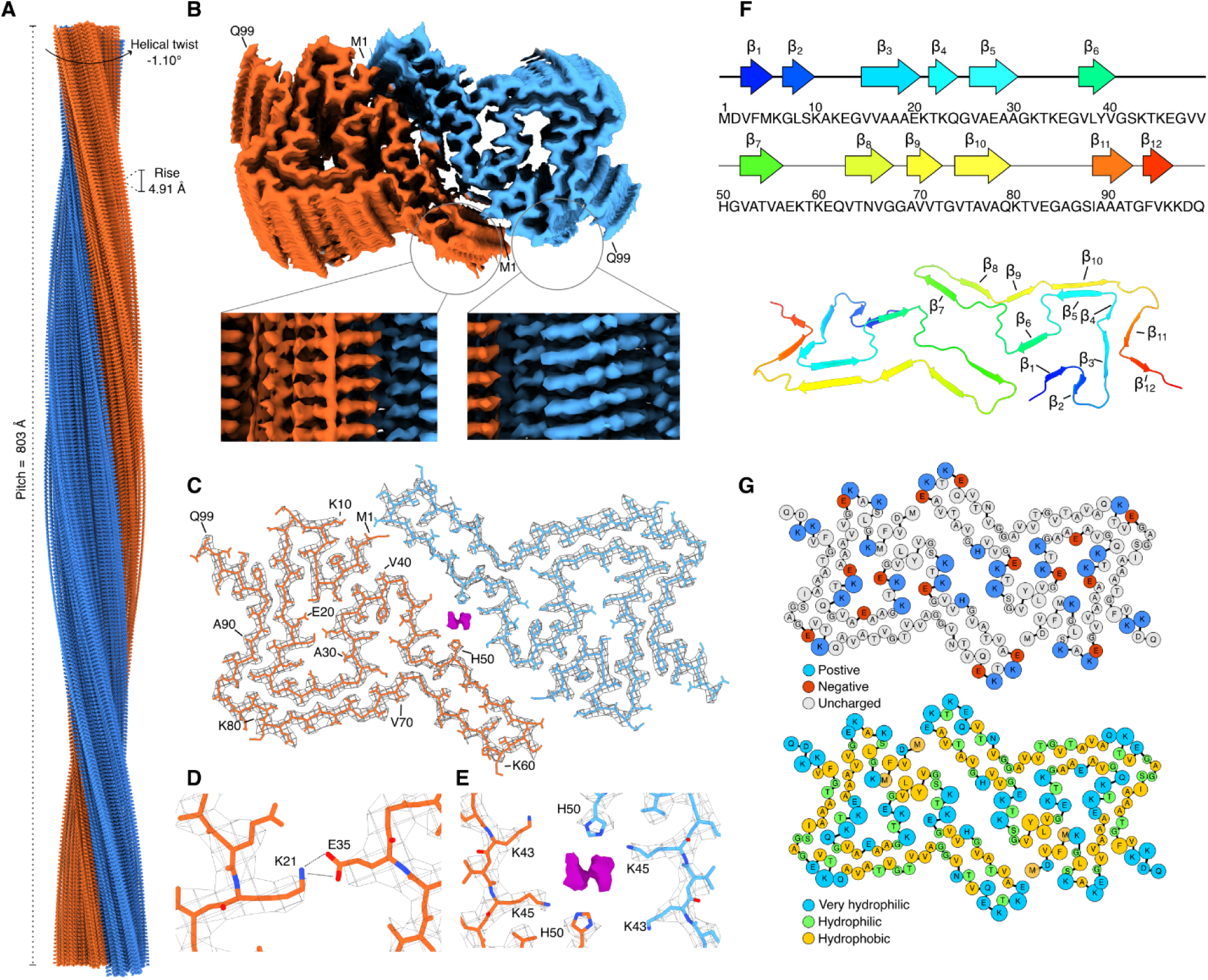
Cryo-EM model of aSyn fibrils formed in aCSF. Side view (A) and top view (B) of the reconstructed Cryo-EM map. Atomic model (residues 1-99) built in the density map (C). The salt bridge (D) formed between K21 and E35 residues. Ligand density embedded inside the fibril core (E). β-sheet arrangements in the structure (F). Charge and hydrophobicity/hydrophilicity distributions (G) in the structure.

### Structural integrity of aCSF fibrils

To understand whether the fibrils formed in aCSF template their structure, a seeded aggregation kinetics experiment was performed (Fig. 4 A) yielding statistically similar endpoint ThT fluorescence intensity values (Supplementary Fig. 8). The residual concentration of soluble (non-aggregated) aSyn was comparatively larger when seeding experiment was done in the aCSF solution (Supplementary Fig. 9). It suggests that the templating process does not produce identical aggregates to the ones used for seeding. Therefore, the resulting aCSF fibrils seeded in PB solution were characterized using Cryo-EM (Fig. 4 B & Supplementary Fig. 3), which showed two distinct fibril conformations (ratio of approx. 1:9). To understand this process further, seeds were resuspended in PB and aCSF solutions (including its variation without specific components or with the addition of 25 µM of EDTA) and imaged using AFM (Fig. 4 C, Supplementary Fig. 10 & 11 A). Based on the visual inspection of fibrils on the mica, the removal of HSA from aCSF fibrils solution or the addition of EDTA affected the aggregate morphology, while removal of other aCSF components showed similar morphologies to the original aCSF fibrils. These fibrils without HSA were mostly shorter that the ones found in the control sample. Nonetheless, such fibrils are not found on the mica once the sample is resuspended back to solution with HSA (aCSF).

**Figure 4.**
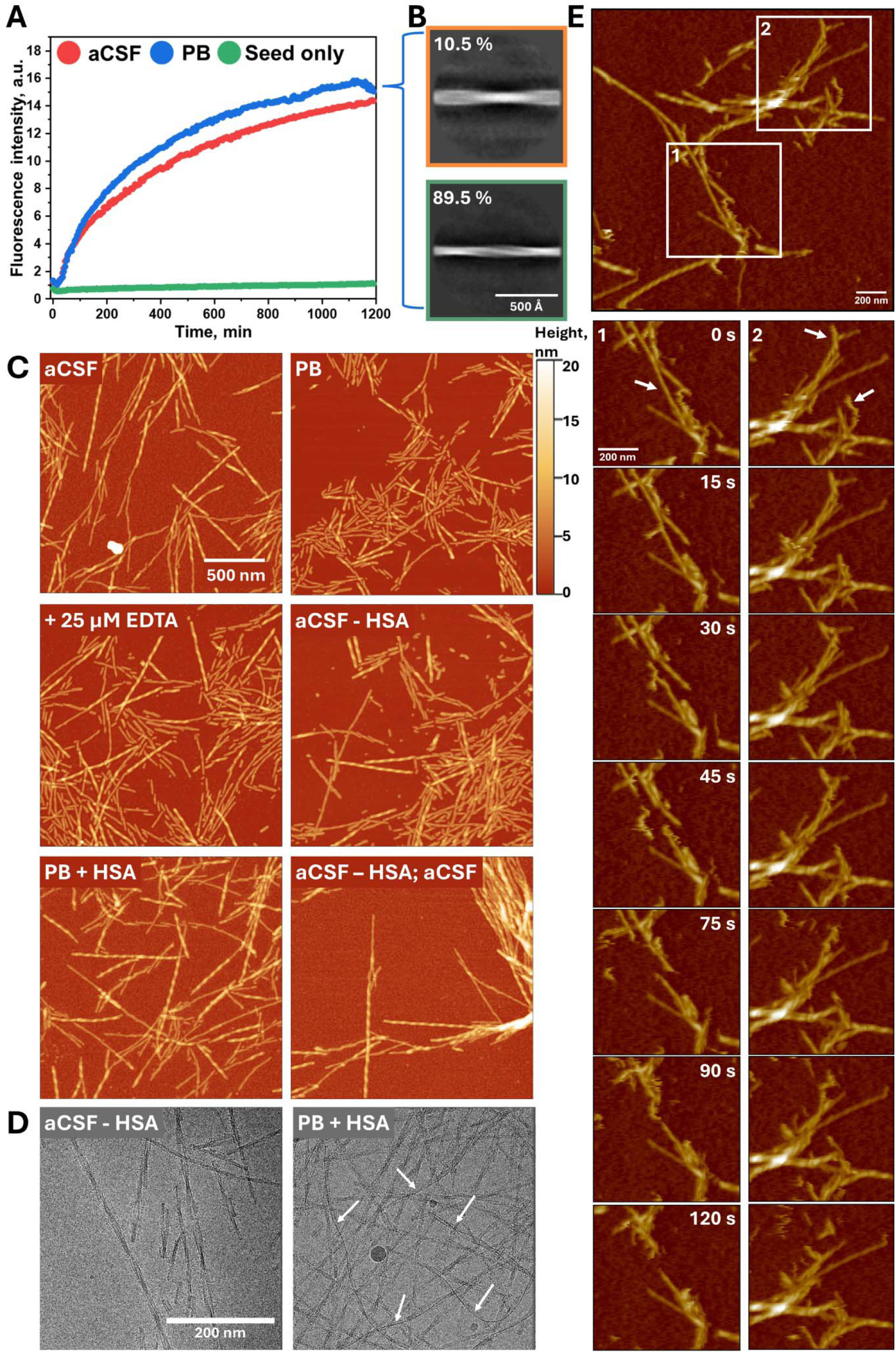
aCSF fibril seeding process in aCSF and PB solutions (A) and the resulting aggregate composition formed in PB solution characterized using Cryo-EM (B). The resulting AFM images (C) of aCSF fibrils 10 minutes after resuspension in different aCSF solution compositions (indicated in top left corner) and Cryo-EM images (D) after incubating the samples for 24 hours. The aCSF – HSA / aCSF indicates the fibrils resuspended in aCSF – HSA solution and then, after 10 minutes incubation resuspended in aCSF. aCSF fibril imaging in PB solution on a mica using HS-AFM (E).

To further investigate the impact of HSA removal from aCSF fibrils solution, we incubated fibrils in the corresponding solutions for 24 hours and performed Cryo-EM (Fig. 4 D & Supplementary Fig. 11 B). Unlike in AFM images, visually observed fibrils of the frozen hydrated micrographs were of similar length. However, the aggregates incubated in the solution with HSA had long thin rod-like structures alongside the fibril’s axis (white arrows), with a couple of cases where two such structures are bound to the fibril, indicating potential HSA binding along the axis of fibrils. Concerning the probable effect of sample preparation (centrifugation, binding on a functionalized mica, freezing), we performed HS-AFM (Fig. 4 E & Supplementary video 1 & 2), where removal of HSA was done after the fibrils were absorbed on the APTES-functionalized mica. The rapid aggregate separation was observed within 10 minutes after the change of solution, while control conditions did not show any significant changes (Supplementary video 3 & 4). These results confirmed the restructurization of aCSF fibrils in HSA-absent environment. However, the excitation-emission matrix (EEM) scans (Supplementary Fig. 12) that can be used to identify different binding mode of ThT to fibrils showed no notable variation in EEM intensity maximum positions throughout the test. This means that even though fibrils restructurize, their binding mode to ThT does not change significantly.

### aSyn fibrils from aCSF exhibit enhanced toxicity

Finally, the sample evaluation on cellular level was conducted (Fig. 5). The SH-SY5Y human neuroblastoma cell line was used to determine whether aSyn aggregation products formed in aCSF or PB solutions possessed distinct effects. The MTT assay was applied for cellular metabolic activity measurement where it indicated potent viability reduction and aggregate type dependent differences (Fig. 5B). Cells treated with aSyn aggregates formed in PB had significantly higher viability compared to ones of aCSF (Supplementary table 4). Here, lower concentrations of fibrils resulted in increased metabolic activity, while the titration of aSyn aggregates formed in aCSF showed an invariable effect. Meanwhile, the LDH release assay revealed that these two types of aggregation products possessed a similar effect on the cell membrane with the most toxic concentration being 1 µM (Fig. 5C). Additionally, cell imaging was conducted to visualise the distribution of aSyn aggregates and possible interaction with SH-SY5Y cells. For such evaluation, the microscopy data was collected before and after cells were washed and media was changed to one without the aggregates (Fig. 5A and Methods section). Despite the equal concentration added, aSyn aggregates formed in aCSF had more active fluorescence signal than the aggregates from PB. After overnight incubation, aSyn aggregates from PB tended to assemble into several individual clusters (Fig. 5A, white arrowheads) while other type of aggregates distributed throughout the cell culture by forming a monolayer. However, the wash-out and media change that potentially had to remove all unbound aggregates from the measurement cell, revealed that aggregation products formed in aCSF had a higher affinity to SH-SY5Y cells compared to fibrils originated in PB.

**Figure 5.**
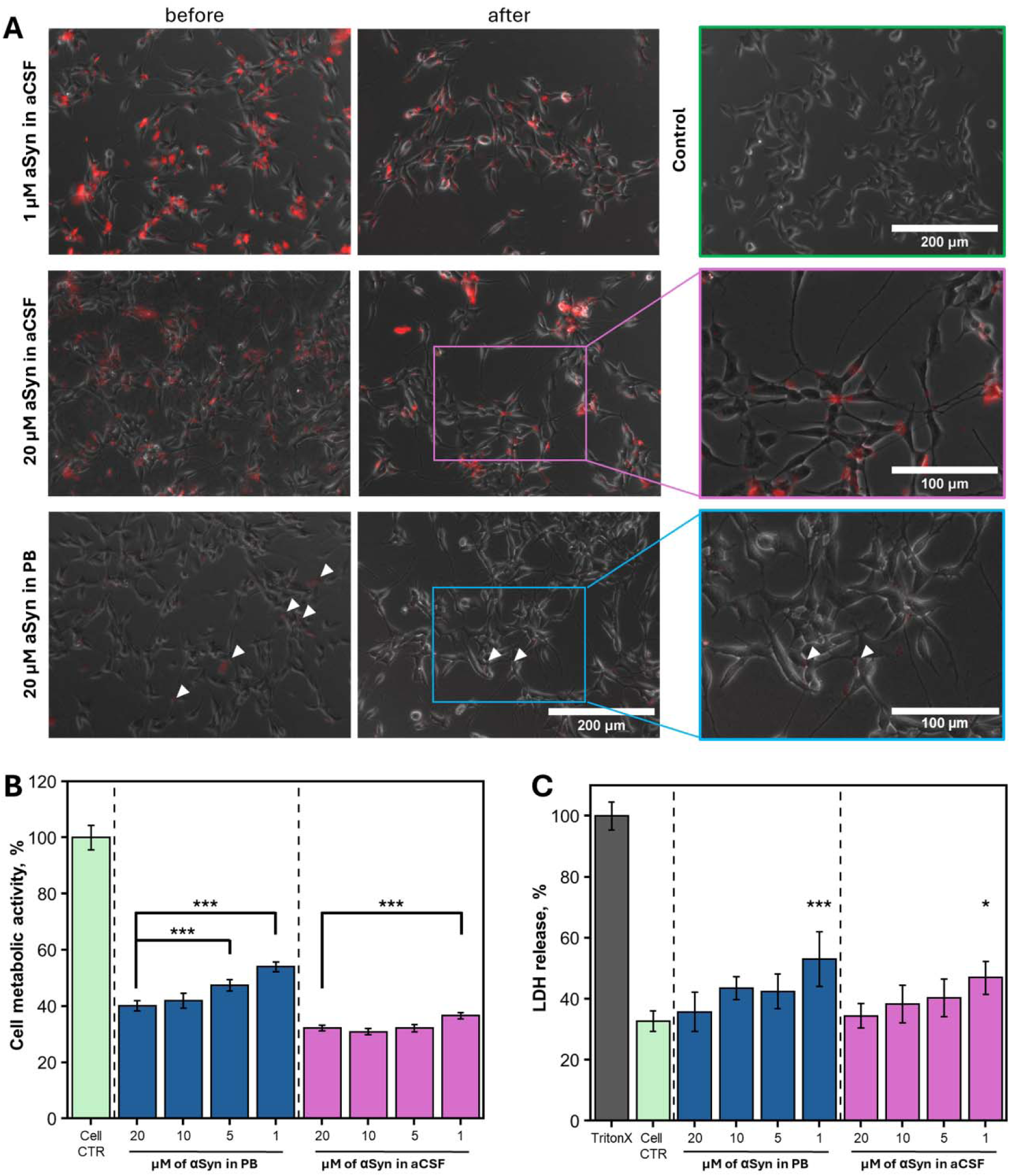
Effect of aSyn aggregates formed in PB or aCSF on SH-SY5Y human neuroblastoma cells. (A) EVOS microscopy images of cells treated with 1 µM and 20 µM of aSyn aggregates formed in aCSF or PB (the aggregation was performed with the labelled aSyn in a ratio of 95% aSyn, 5% mCherry-aSyn). Untreated SH-SY5Y cells were used as control. Imageswere collected before and after media change. Changes in cell metabolic activity (B) and cytotoxicity (C) after incubation with different types of aSyn aggregates in concentration dependent manner. One-way ANOVA Bonferroni means comparison for each condition was conducted to determine differences between sample concentrations (\**p* < 0.05, \*\**p* < 0.01, \*\*\**p* < 0.001). For LDH release, the comparison was with cell control.

## Discussion

Growing knowledge in the field of synucleinopathies, including Parkinson’s disease, has indicated the role of aSyn aggregation as a critical driver of disease progression and heterogeneity^61^. A key focus has been on the protein’s properties to adopt distinct conformations and propagate in a prion-like manner. However, recent evidence^62–65^ shows that aSyn extracted from brain tissue or cerebrospinal fluid fails to template the recombinant protein *in vitro*. These findings shift the research to a new direction, where the native aggregation environment in the brain tissue, cells or CSF may play a significant role in determining fibril formation, structure and morphology. These differences raise the importance of studying aSyn aggregation in physiologically relevant conditions. To address this point of view, we used artificial cerebrospinal fluid to recreate the environment closer to *in vivo* conditions where aSyn aggregates formation and templating processes occur.

Our findings demonstrate that components from artificial cerebrospinal fluid dictate the formation of a specific aSyn fibril structure. As expected, divalent metal ions (Ca^2+^ and Mg^2+^) emerged as key modulators of protein aggregation kinetics and structural variation observed using FTIR. While calcium and magnesium are well known to interact with negatively charged C-terminal region of aSyn^66,67^, the aCSF formulation includes HSA that chelates Me^2+^ and is proven to increase its interaction with aSyn N- and C-terminals once Cu^2+^ ions are bound to it^68,69^. It is probable that such interaction specifically occurs in our case, as the removal of HSA or the addition of EDTA (to bind all divalent metals) resulted in the fragmentation of preformed aCSF fibrils and failure to 100% replicate its conformation. This suggests that fibrils produced in aCSF are dynamic when the critical components that stabilize their structure are removed, which contradicts the common belief that once fibrils form, they retain a fixed structure due to its energy minimum^31^. Such fibril morphological instability complicates matters for *in vitro* research due to sample preparation limitations (fibril resuspension in measurement buffer solutions, extraction method from *ex vivo* samples, desalting procedures, PMCA and many other factors)^26,40,62,70,71^. It could be that *ex vivo* fibrils transform, resulting in a different morphology once observed using AFM, Cryo-EM or other methods. This environment-specific shift may also explain why it is not possible to replicate the extracted fibrils *in vitro*^62–65^.

We also determined the Cryo-EM structure of newly formed aCSF fibrils which does not have an exact match with any currently known resolved structure. However, its structural assembly resembles electron density pocket of aggregates found in multiple system atrophy (MSA) patients^63^. Although we show symmetrical fibril core that differs from solved *ex vivo* MSA type 1 and type 2 aggregates that have asymmetrical conformations, the electron density pocket in all mentioned cases is co-ordinated with identical AA positions (K43, K45, H50)^41,63,72^. Additionally, there are other cases where lysine and histidine are shown to coordinate the ligand at the side of the aggregate^42^ or AAs tend to interact with each other within the same protofilament (inter-rung connection between H50 and K45)^73^, meaning the molecule that coordinates these amino acids yields a distinct aggregate formation pathway. In our case, the probable molecules that may fit the pocket are glucose, glutamine and metal ion complexes. However, more research needs to be done in order to find this target substance, especially, when the instability of fibrils complicates their extraction from the solution, thus limiting the likelihood of determining the ligand.

Another important matter to consider is the aSyn aggregate effect on neuroblastoma cells. aCSF fibrils tended to adhere to the cell surface significantly more than those of a control group (PB). There is evidence that specific ions such as calcium or zinc, found in cerebrospinal fluid, alter aSyn fibril morphology and promote different aggregation pathways. These fibrils exhibit adhesive properties to cell membrane and internal structures, such as pre-synaptic vesicles^32,67,74^, where Me^2+^ concentration fluctuation may occur^75,76^. In fact, our Cryo-EM data shows that outer surface of the fibrils carries positive charged areas, which can be attracted to a negatively charged cell membrane^77^. Moreover, structural features such as hydrophobic pockets and β-sheet stacking could influence their interaction with cell membranes^39,65^, enhancing affinity to the lipid bilayer or cellular components. This may lead to extensive cellular internalization, membrane disruption or oxidative stress^78–80^. Additionally, the cell toxicity assay revealed an interesting tendency. While cell metabolic activity was reduced at higher aggregate concentrations, the LDH release that mostly describes damage to the cell wall was significantly higher when lower aggregate concentration was added. It could be that aSyn fibrils tend to dissociate to smaller fragments at lower concentrations due to excess amounts of cell media components or being prone to mechanistic stress (cell movement, pipetting)^81,82^. Separated fibrils, albeit their strong affinity towards each other, have less probability to interact (to clump together), resulting in binding directly to the cell membrane and damaging it as smaller, more toxic fragments^83^.

These findings provide evidence that the molecular and ionic interfaces of cerebrospinal fluid play an important role in fibril morphology, suggesting that fibril structure is environment-dependent rather than dictated by properties of aSyn secondary structure. Recreatingphysiological conditions for aggregate extraction and templating may be the key factor enabling our further understanding about neurodegenerative diseases. Furthermore, ability to seed identical aggregate structures could contribute to creating kits for detection of disease-specific biomarkers that are currently are not efficient enough^84^.

## Conclusion

In conclusion, artificial cerebrospinal fluid has shifted aSyn aggregation towards an environment-specific conformation that was solved using Cryo-EM (2.8 Å resolution). This fibril conformation has not previously been observed in any aSyn-related studies. The main contributors to this effect were found to be calcium, magnesium ions and human serum albumin. The latter proved to be the main factor in stabilizing the fibril conformation within the aCSF environment. Moreover, the aCSF fibrils exhibited increased affinity to SH-SY5Y neuroblastoma cells.

## Acknowledgements

We would like to thank Michael Adams and Nadishka Jayawardena from Thermo Fisher Center for Electron Microscopy – Eindhoven NanoPort and for helping to record the Cryo-EM data in their facility. We also thank Jiří Nováček and Zuzana Hlavenková from CEITEC for supporting in measuring additional data in their facility via Instruct-ERIC proposal. Finally, we are also grateful to Sjors Scheres from the MRC Laboratory of Molecular Biology for suggestions and comments on generating an initial Cryo-EM map of aCSF amyloid fibrils.

## Author Contribution

D.S.: Conceptualization, Methodology, Validation, Formal analysis, Investigation, Writing - Original Draft, Writing - Review & Editing, Supervision, Funding Acquisition. A.S.: Conceptualization, Methodology, Validation, Formal analysis, Investigation, Data Curation, Writing - Original Draft, Writing - Review & Editing, Visualization, Supervision, Project Administration. R.S.: Methodology, Formal Analysis, Investigation, Writing - Original Draft, Writing - Review & Editing, Visualization. R.T.: Methodology, Investigation. M.Z.: Methodology, Validation, Formal Analysis, Investigation, Writing - Review & Editing. A.J.: Investigation. U.V.: Investigation, Writing - Review & Editing. V.S.: Resources, Writing - Review & Editing, Funding Acquisition.

## Data and materials availability

All data needed to evaluate the conclusions in the paper are present in the paper and/or Supplementary Materials. The raw data used in this paper have been tabulated and are available on Mendeley Data: (10.17632/d3cby7cv57.1). The reconstructed cryo-EM map was deposited in the Electron Microscopy Data Bank with the accession codes EMD-52833 The coordinates of the fitted atomic model were deposited in the Protein Data Bank under the accession code 9IC7. All other relevant data are available from the corresponding author upon reasonable request.

