## Supplementary materials for "Formation of Condition-Dependent Alpha-Synuclein Fibril Strain in Artificial Cerebrospinal Fluid"

Table 1. Composition of aCSF.

|  | **Concentration in 10x solution, mM** | | | | | | | | | **1x, mM** |
| --- | --- | --- | --- | --- | --- | --- | --- | --- | --- | --- |
| ***Component*** | **PB (A)** | ***B*** | ***C*** | ***D*** | ***E*** | ***F*** | ***G*** | ***H*** | ***I*** | **aCSF** |
| NaCl | 1270 |  |  |  |  |  |  |  |  | 127 |
| KCl | 18 |  |  |  |  |  |  |  |  | 1.8 |
| Na_2_HPO_4_ | 78.1 |  |  |  |  |  |  |  |  | 7.81 |
| NaH_2_PO_4_ | 31.9 |  |  |  |  |  |  |  |  | 3.19 |
| KH_2_PO_4_ | 12 |  |  |  |  |  |  |  |  | 1.2 |
| Urea |  | 6.5 |  |  |  |  |  |  |  | 0.65 |
| L-glutamine |  |  | 7 |  |  |  |  |  |  | 0.7 |
| HSA |  |  |  | 0.0615 |  |  |  |  |  | 0.00615 |
| Cholesterol |  |  |  |  | 0.052 |  |  |  |  | 0.0052 |
| Sodium lactate |  |  |  |  |  | 24 |  |  |  | 2.4 |
| CaCl_2_ |  |  |  |  |  |  | 14 |  |  | 1.4 |
| MgCl_2_ |  |  |  |  |  |  |  | 13 |  | 1.3 |
| Glucose |  |  |  |  |  |  |  |  | 40 | 4 |

Table 2. Experimental design of aggregation study. The grey box marks the missing components from the final reaction mixture.

|  | | **PB (A)** | ***B*** | ***C*** | ***D*** | ***E*** | ***F*** | ***G*** | ***H*** | ***I*** | ***aSyn*** | ***ThT*** |
| --- | --- | --- | --- | --- | --- | --- | --- | --- | --- | --- | --- | --- |
| Reaction mixtures | aCSF |  |  |  |  |  |  |  |  |  |  |  |
|  | aCSF - urea |  |  |  |  |  |  |  |  |  |  |  |
|  | aCSF - glutamine |  |  |  |  |  |  |  |  |  |  |  |
|  | aCSF - HSA |  |  |  |  |  |  |  |  |  |  |  |
|  | aCSF - cholesterol |  |  |  |  |  |  |  |  |  |  |  |
|  | aCSF - sodium lactate |  |  |  |  |  |  |  |  |  |  |  |
|  | aCSF – CaCl_2_ |  |  |  |  |  |  |  |  |  |  |  |
|  | aCSF – MgCl_2_ |  |  |  |  |  |  |  |  |  |  |  |
|  | aCSF - glucose |  |  |  |  |  |  |  |  |  |  |  |
|  | ***PB*** |  |  |  |  |  |  |  |  |  |  |  |
|  | ***PB*** + glucose |  |  |  |  |  |  |  |  |  |  |  |
|  | ***PB*** + MgCl_2_ |  |  |  |  |  |  |  |  |  |  |  |
|  | ***PB*** + CaCl_2_ |  |  |  |  |  |  |  |  |  |  |  |
|  | ***PB*** + sodium lactate |  |  |  |  |  |  |  |  |  |  |  |
|  | ***PB*** + cholesterol |  |  |  |  |  |  |  |  |  |  |  |
|  | ***PB*** + HSA |  |  |  |  |  |  |  |  |  |  |  |
|  | ***PB*** + glutamine |  |  |  |  |  |  |  |  |  |  |  |
|  | ***PB*** + urea |  |  |  |  |  |  |  |  |  |  |  |

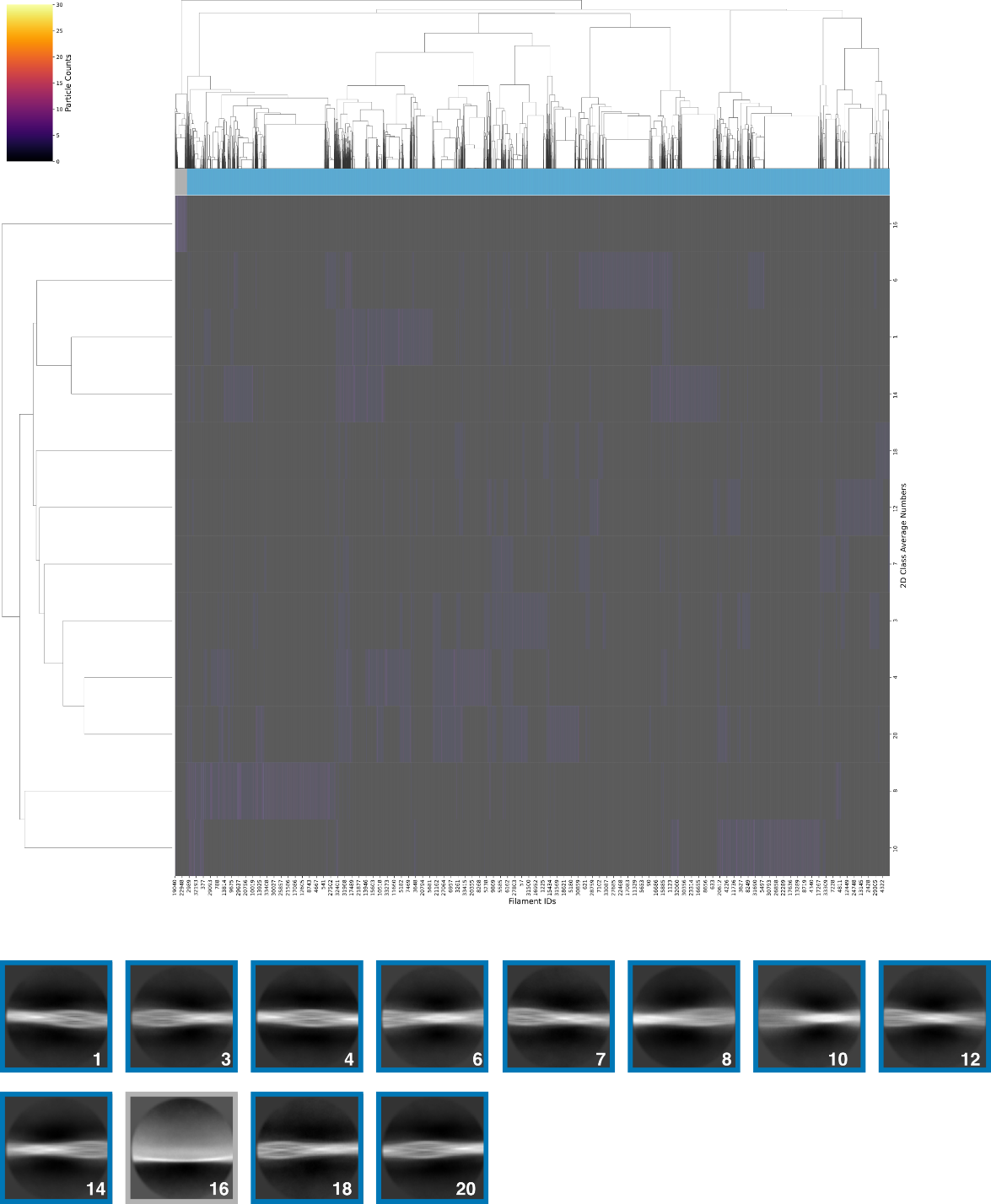

Figure 1. aCSF fibril distribution.

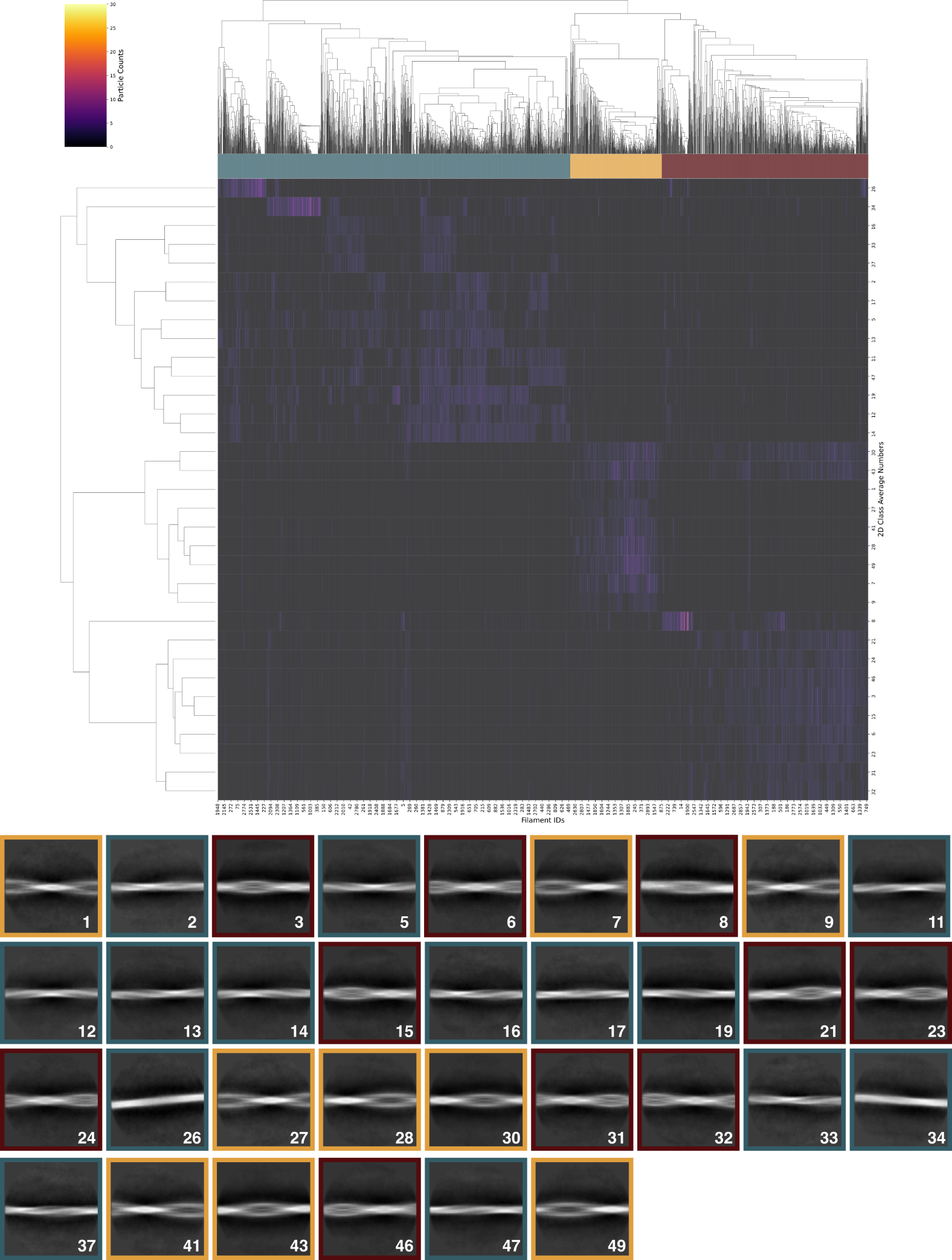

Figure 2. PB fibril distribution.

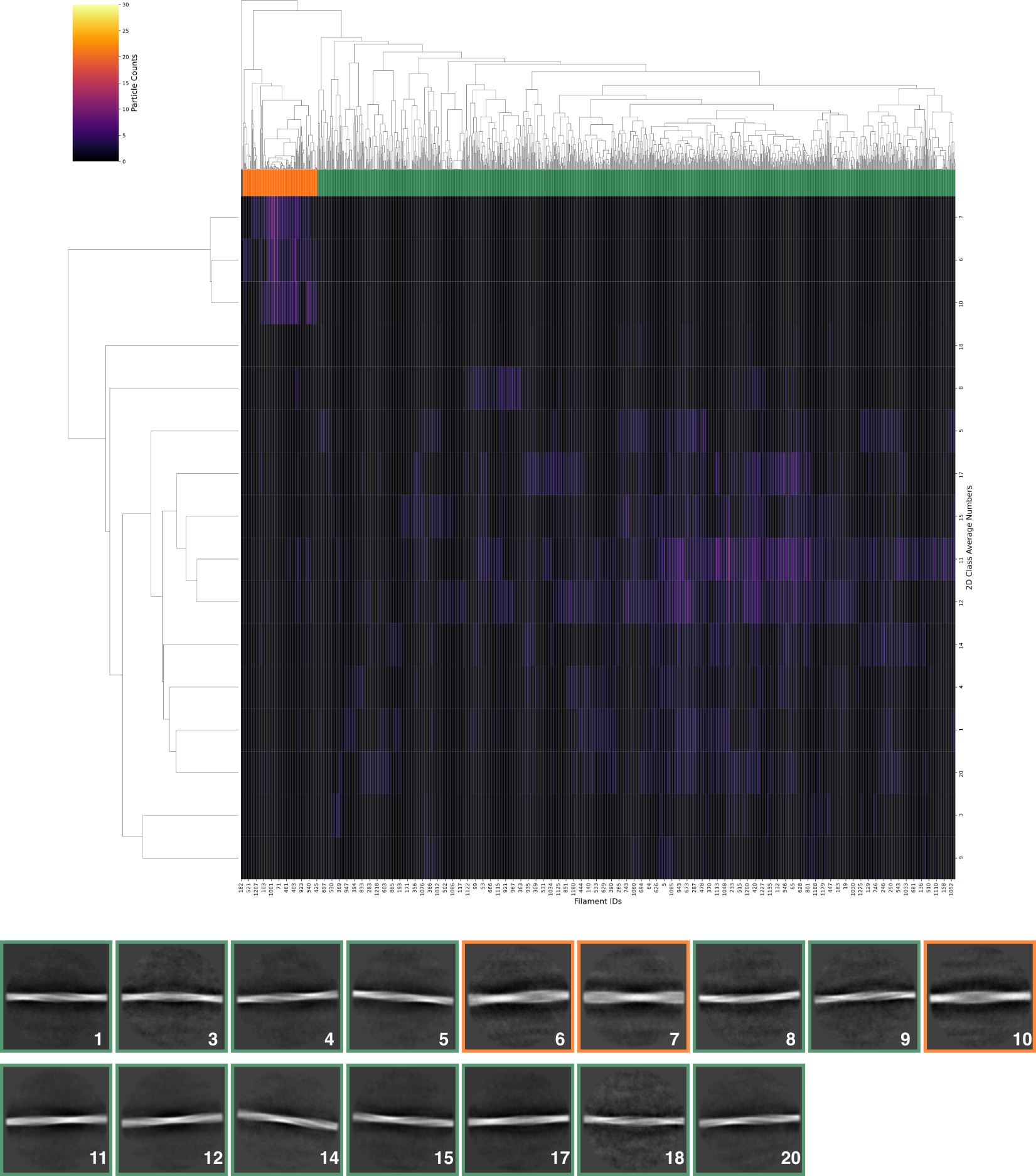

Figure 3. Distribution of aCSF fibrils seeded in PB.

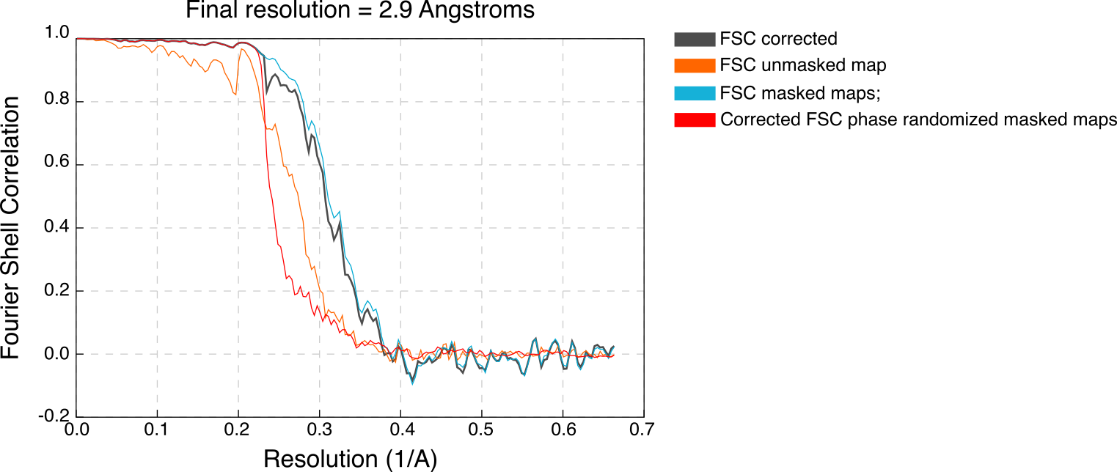

Figure 4. Fourier shell correlation (FSC) curves after final post-processing in Relion.

Table 3. Cryo-EM data collection and modelling statistics.

|  | Alpha-synuclein in aCSF | Alpha-Synuclein in PBS | Asyn_ascf_ seeded to PB conditions |
| --- | --- | --- | --- |
| **Data collection** |  |  |  |
| Magnification | 165000 | 150000 | 92000 |
| Pixel size (Å) | 0.754 | 0.95 | 1.1 |
| Defocus range (μm) | -2.0 to -1.0 | -2.2 to -1.2 | -2.2 to -1.2 |
| Voltage (kV) | 300 | 200 | 200 |
| Camera | Falcon 4i | Falcon 4i | Falcon 3CE |
| Microscope | Krios | Glacios 2 | Glacios |
| Energy filter slit width (eV) | 10 | - | - |
| Exposure time | 3.37 | 3.99 | 46.33 |
| Number of eer fractions | 30 | 40 | 30 |
| Total dose (e^-^/Å^2^) | 22.65 | 30 | 30 |
| **Data processing** |  |  |  |
| Micrographs | 3510 | 3199 | 857 |
| Picked particles | 161452 | 6516 | 1014 |
| Box size (pixel) | 384 | 384 or 1024 downscaled to 256 | 1024 downscaled to 256 |
| Inter-box distance (Å) | 18.9 | 30/20 | 17.27 |
| Segments extracted | 1253042 | 122335/33022 | 6750 |
| Segments used for 3D final reconstruction | 20772 | - | - |
| Helical twist (º) | -1.11 | - | - |
| Helical rise (Å) | 4.91 | - | - |
| Symmetry imposed | C2 | - | - |
| Map resolution FSC 0.143 (Å) | 2.9 | - | - |
| **Refinement** |  |  |  |
| Model resolution (Å)  FSC threshold  Map sharpening factor (Å2) | 2.9  0.143  -42.2007 | - | - |
| **Model** **composition** |  |  |  |
| Non-hydrogen atoms  Protein residues  Ligands | 9688  1386  0 | - | - |
| B factors (Å^2^) min/max/mean | 14.10/96.52/48.71 |  | - |
| **R.m.s.** **deviations** |  |  |  |
| Bond lengths (Å)  Bond angles (°) | 0.003  0.504 | - | - |
| **Validation** |  |  |  |
| MolProbity score  Clashscore  Rotamer outliers (%) | 2.23  6.02  4.35 | - | - |
| **Ramachandran plot** |  |  |  |
| Favored (%)  Allowed (%)  Disallowed (%) | 0  6.19  93.81 | - | - |

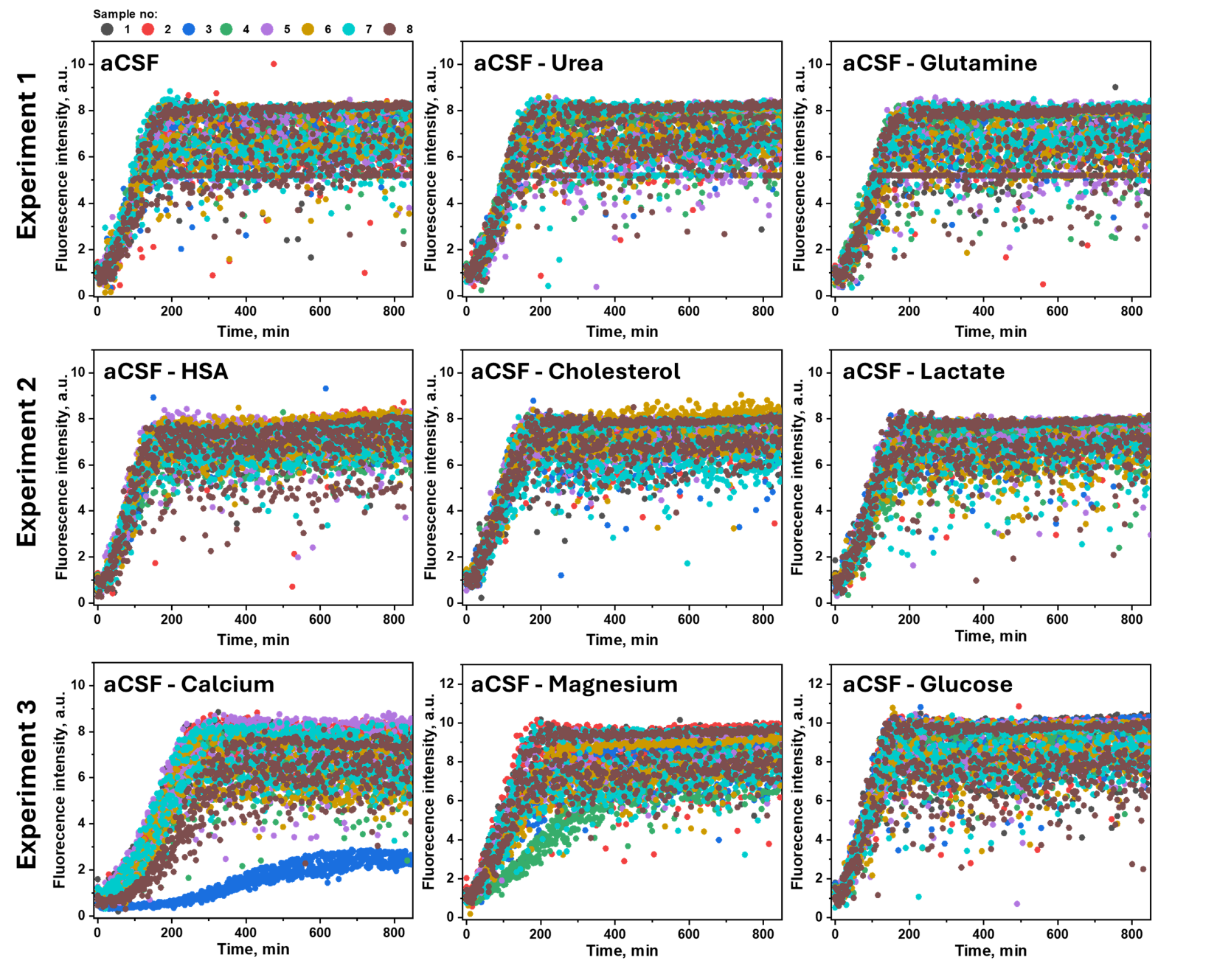

Figure 5. Seeded aggregation kinetics of aSyn. After de novo aggregation, aggregates formed in each condition were used to seed the monomer aSyn (10% of seeds added). For data measurements (experiments 1 - 3), enhanced dynamic range was used, disallowing the fluorescence intensity comparison due to automatic fluorescence intensity adjustments.

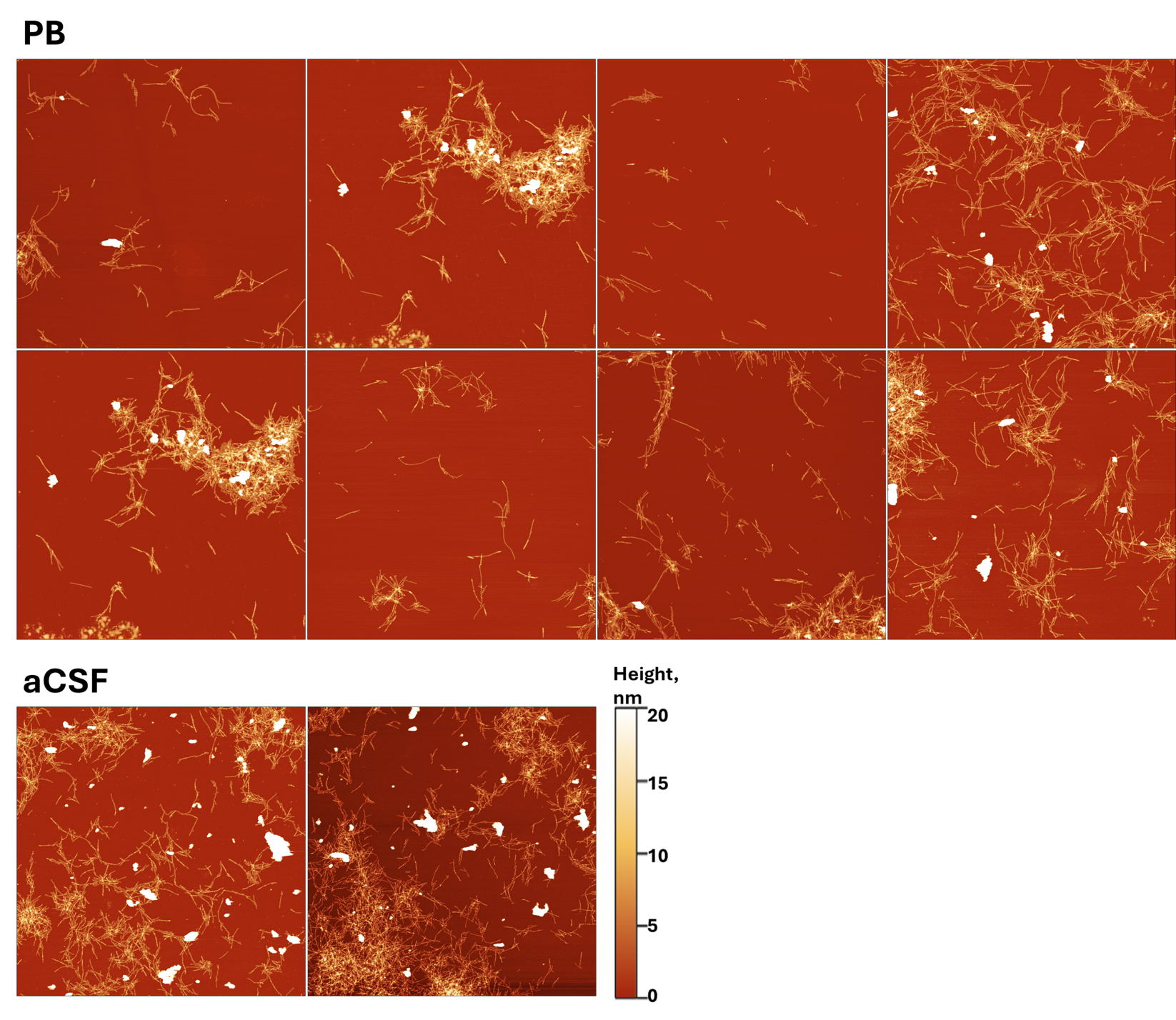

Figure 6. PB ir aCSF fibrils

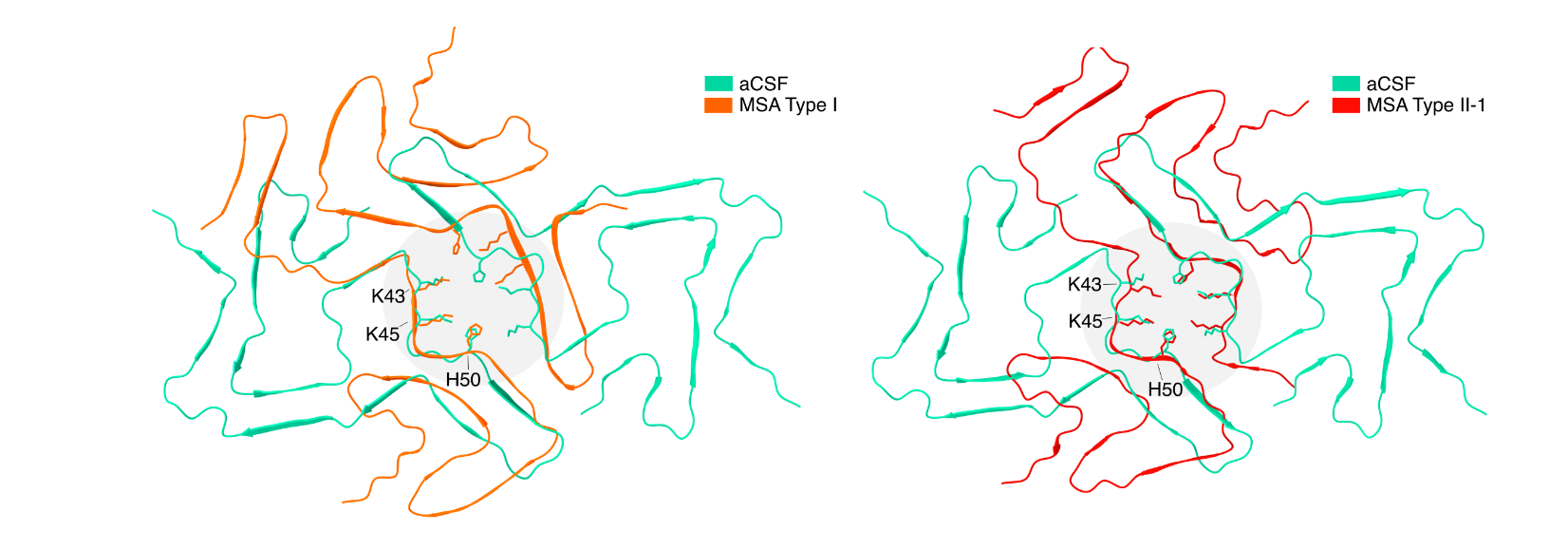

Figure 7. Cryo-EM model comparison between aggregates formed in aCSF and found in patients from MSA (MSA Type I (PDB id: 6XYO) and MSA Type II-1 (PDB id: 6XYP)).

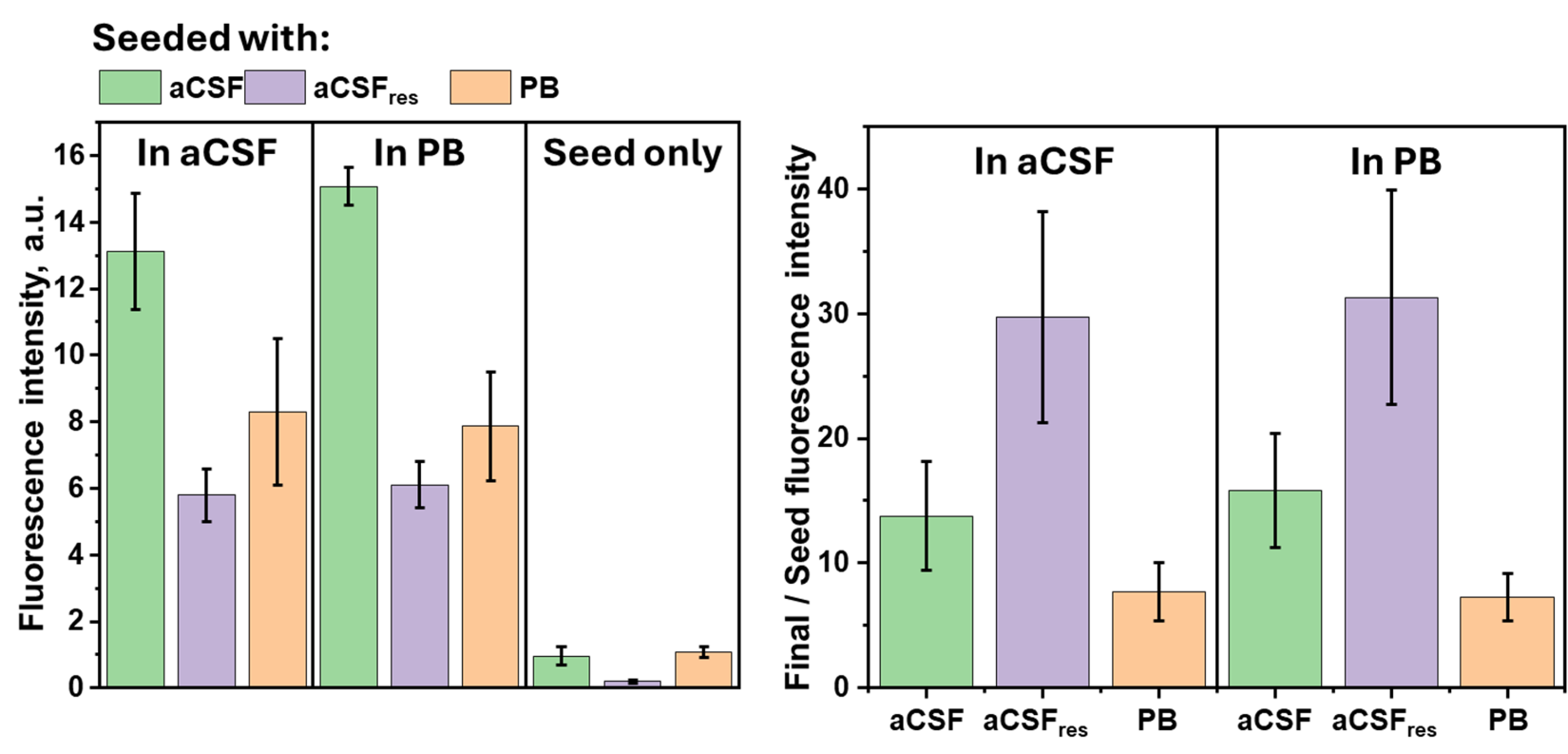

Figure 8. Seeded aggregation kinetics endpoint ThT fluorescence intensity values and ratio between final and seed fluorescence intensities. aCSF_res_ - resuspended aCSF fibrils with PB and incubated (7 days).

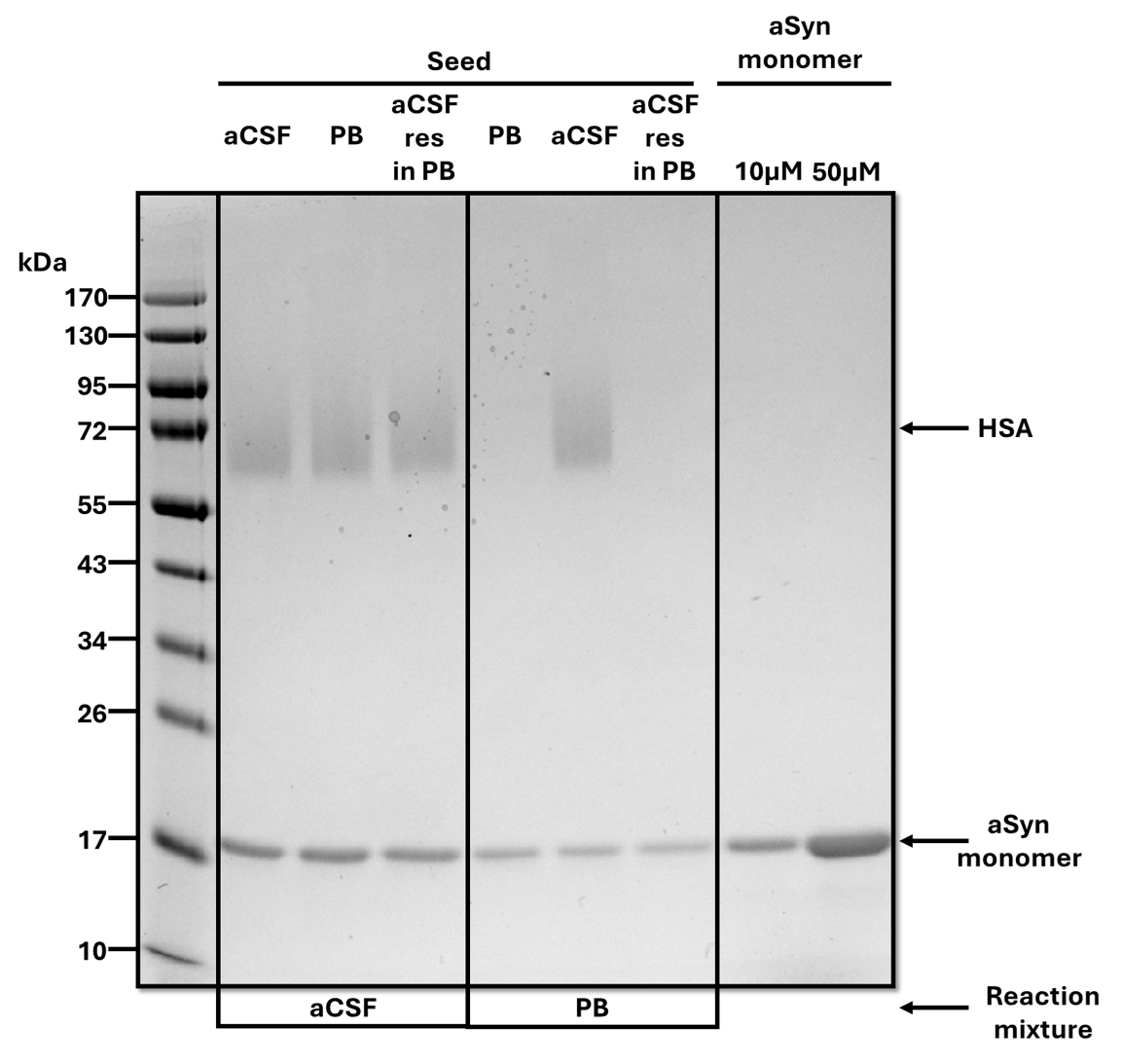

Figure 9. SDS-page of final aggregation of seeded sample supernatant.

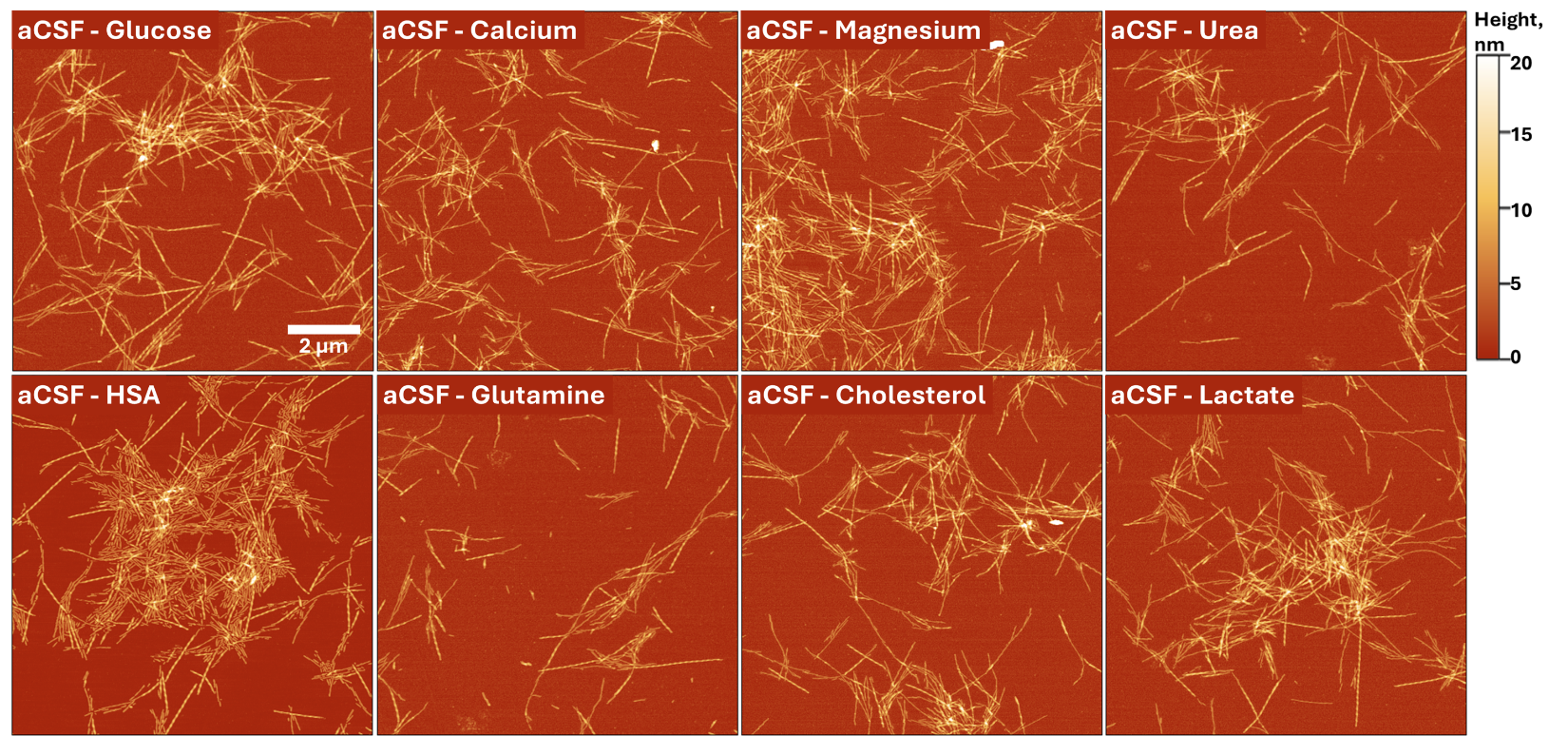

Figure 10. AFM images of aCSF fibrils resuspended in the solution with one of its components removed (indicated top left corner of each picture).

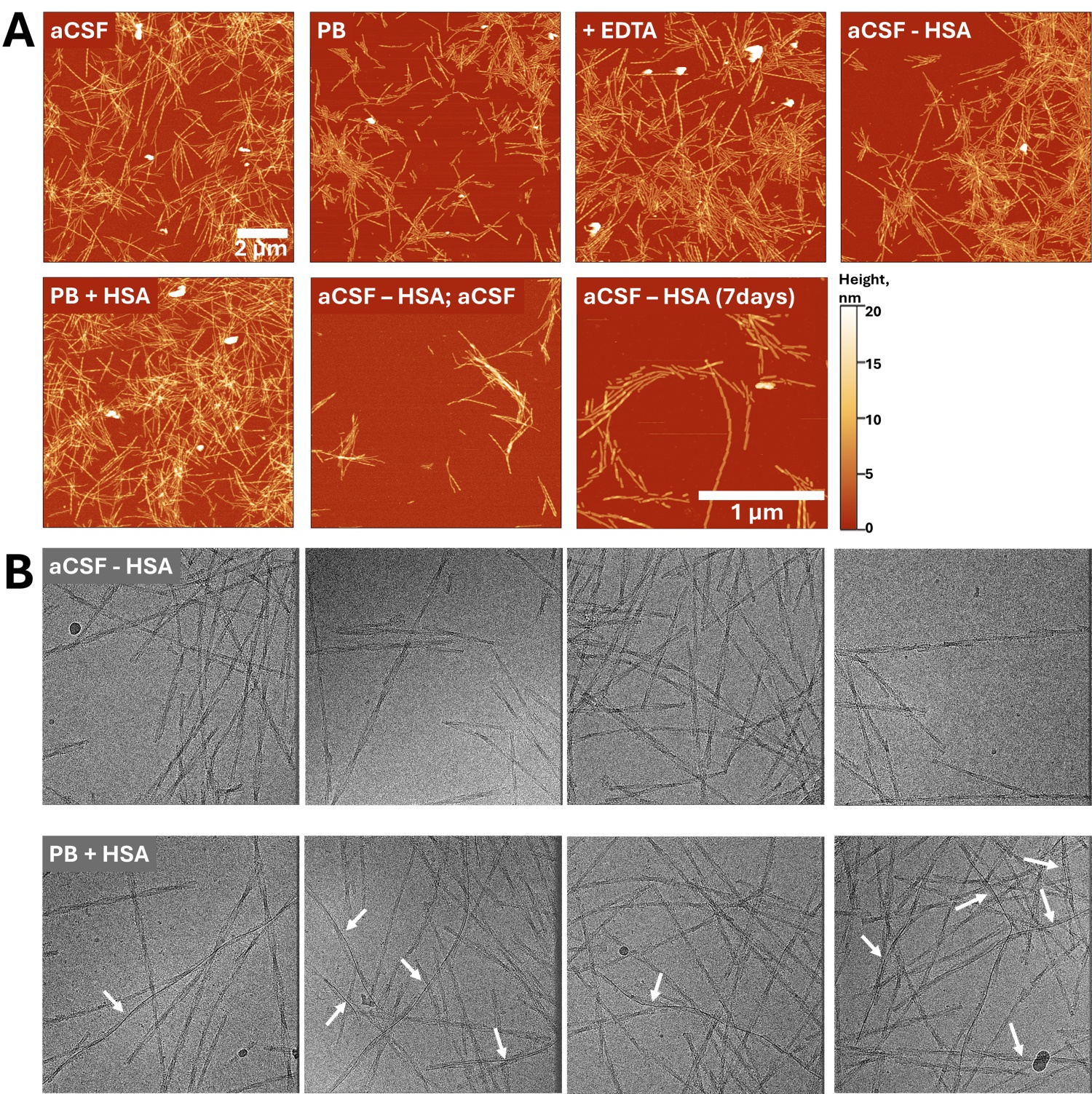

Figure 11. AFM (A) and Cryo-EM (B) images of aCSF fibrils resuspended in different aCSF compositions. In case of aCSF – HAS; aCSF, fibrils were resuspended in aCSF – HSA and then resuspended in aCSF.

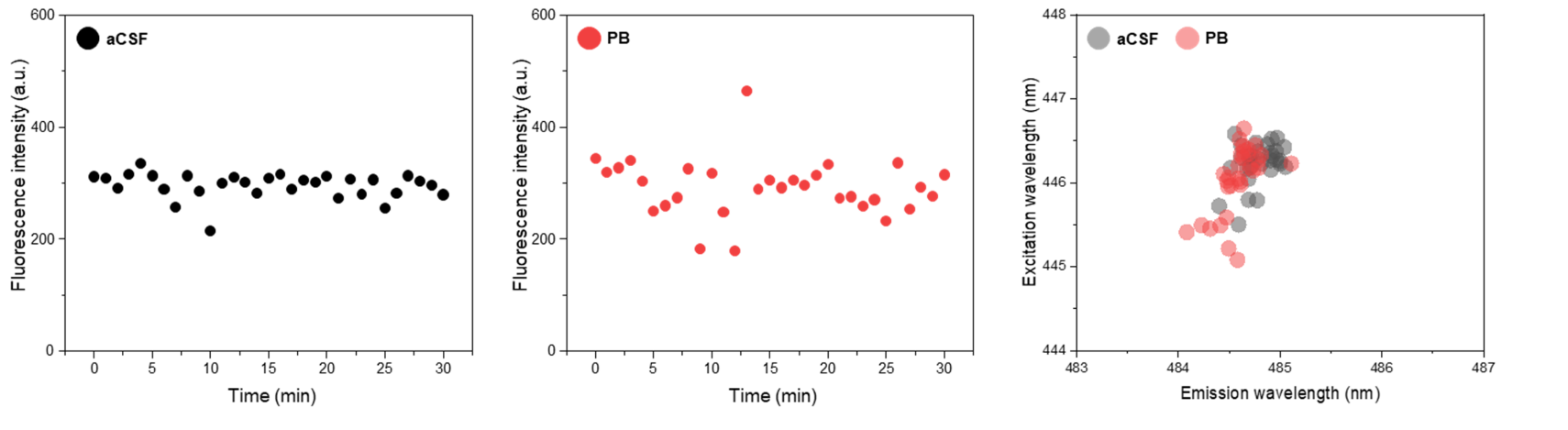

Figure 11. EEM intensity maximum position variation over time of aCSF fibril-bound ThT once they are resuspended in aCSF or PB solutions.

Table 4. ANOVA means comparison of aSyn aggregates effect to cell. The comparison is done between equal concentration of fibrils that were produced at different conditions (aCSF and PB). “NS” corresponds to not significant.

|  | *Significance* | |
| --- | --- | --- |
| **aSyn concentration (µM)** | **MTT** | **LDH** |
| 20 | 0.001 | NS |
| 10 | 0.001 | NS |
| 5 | 0.001 | NS |
| 1 | 0.001 | NS |
